# Virtual screening identifies novel high-affinity σ_1_ receptor ligands

**DOI:** 10.1101/699793

**Authors:** Daniel A. Greenfield, Hayden R. Schmidt, Piotr Sliz, Andrew C. Kruse

## Abstract

The σ_1_ receptor is a transmembrane protein implicated in several pathophysiological conditions, including neurodegenerative disease^1^, drug addiction^2^, cancer^3^, and pain^4^. However, there are no high-throughput functional assays for σ_1_ receptor drug discovery. Here, we assessed high-throughput structure-based computational docking for discovery of novel ligands of the σ_1_ receptor. We screened a library of over 6 million compounds using the Schrödinger Glide package, followed by experimental characterization of top-scoring candidates. 77% of tested candidates bound σ1 with high affinity (10-550 nM). These include compounds with high selectivity for the σ_1_ receptor compared to the genetically unrelated but pharmacologically similar σ_2_ receptor, as well as compounds with substantial cross-reactivity between the two receptors. These results establish structure-based virtual screening as a highly effective platform for σ_1_ receptor ligand discovery.

## Main text

The σ_1_ receptor is an enigmatic intracellular membrane protein that is an active drug target for the treatment of myriad conditions including neurodegenerative disease^1^, pain^4^, cancer^3^, and drug abuse^2^. The receptor binds a chemically diverse set of ligands with high affinity, including many compounds that are designed to bind to other receptors, such as antidepressants, antipsychotics, and NMDA receptor ligands, among others^5^. Ligands targeting the σ_1_ receptor are in clinical trials for treatment of neuropathic pain^6^, ischemic stroke^7^, and Alzheimer’s disease^8^. The σ_1_ receptor is expressed throughout the human body but is most highly expressed in the nervous system^5^. It is genetically unique, with no appreciable sequence similarity to any other gene in the human genome^5^. Even the pharmacologically similar σ_2_ receptor is unrelated in sequence, despite its similar ligand affinity profile^9,10^.

Despite decades of study, structural data for the σ_1_ receptor have only recently become available^11^. The receptor crystallized as a homotrimer, though it is known to exist in multiple oligomeric states^12-12^. Each monomer has an occluded ligand binding site containing Glu172 and Asp126^11^, which are essential for ligand binding^15^. The first crystal structure of the human σ_1_ receptor was solved with the receptor in complex with PD144418 at 2.5 Å resolution (PDB: 5HK1). Four other receptor-ligand complexes with ligands 4-IBP, haloperidol, NE-100, and (+)-pentazocine have been reported at crystallographic resolutions of 2.8-3.1 Å^11,16^. As is typical of σ_1_ receptor ligands, these co-crystallized ligands are structurally divergent from one another. Nonetheless, all five σ_1_ receptor-ligand complexes solved to date share some key features, the most notable of which is an essential electrostatic interaction between a basic amine moiety in the ligand and the side chain of receptor residue Glu172^11,16^. Given this constraint, we hypothesized that the σ_1_ receptor would be an ideal candidate for *in silico* ligand screening.

*In silico* screening uses software to estimate ligand binding energy, enabling efficient prioritization of compounds while sampling a larger chemical space than is readily accessible to experimental methods^17^. Schrödinger’s Drug Discovery Platform Glide module^18,19^ is a widely used ligand docking software and is used as the preferred docking system by the structural biology groups affiliated with SBGrid^20^. Glide converges upon optimally docked compounds by sampling the ligand in various states and orientations within the user-defined binding site^18,19^. Candidate ligands that pass through initial space screens (*i.e*., those that are sterically capable of fitting into the target site), are scored at increasing precision levels until the final Glide Score is calculated, with lower Glide Scores indicating a higher predicted binding affinity^19^. Glide docking has three precision levels, High Throughput Virtual Screening (HTVS), Standard Precision (SP), and Extra Precision (XP). HTVS is a less stringent method that eliminates ligands from the sample pool if their volume exceeds the binding site volume. SP and XP are slower, more intensive algorithms that predict binding chemistry. In the case of the σ_1_ receptor, where no high-throughput biological assays are available, computational docking is potentially useful for identifying novel compounds that are chemically dissimilar to both one another and to known σ1 receptor ligands.

Prior to docking experiments, a grid for Glide docking was established as a 10 Å cube centered between the carboxylates of Glu172 and Asp126. In pilot docking test, this resulted in known binders occupying a similar pose within the orthosteric site to that observed in crystal structures. Enrichment plots were calculated to assess docking paradigms, following previously reported approaches^17,21^. To compare enrichment with different approaches, we used 32 conformers of 15 known binders and 100,000 randomly selected eMolecules compounds of similar molecular weight into a test subset (see Methods). The enrichment plots generated over this subset confirmed that we were adequately predicting the binding mode of σ_1_ ligands in the docking protocol. The docking algorithm recovered (scored) over 90% of known binders, with a high proportion of the known binders being in the top 10% of scored compounds. Additionally, through visual confirmation of Glide successfully docking the co-crystallized ligand in a similar configuration as it is bound (Figure 2A), the docking protocol demonstrated potential predictive value.

**Figure 1.**
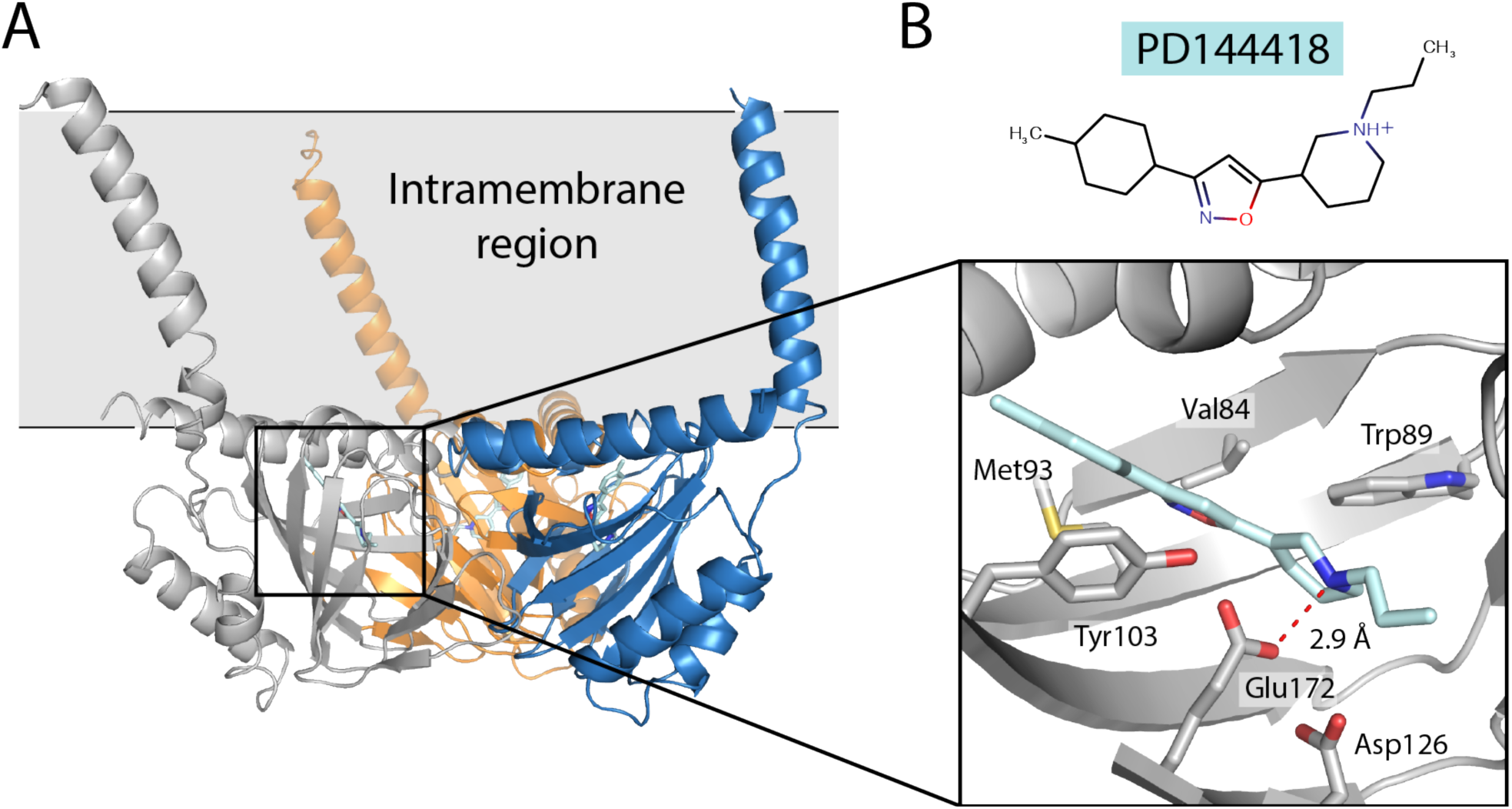
Structure of the human σ_1_ receptor bound to PD144418 (PDB: 5HK1). (A) Overall architecture of the human σ_1_ receptor. The receptor crystallized as a homotrimer, with each monomer binding one ligand molecule in an occluded binding pocket. (B) Close-up of the σ_1_ receptor’s ligand binding pocket with several amino acid side chains depicted as sticks and PD144418 shown in cyan.

**Figure 2.**
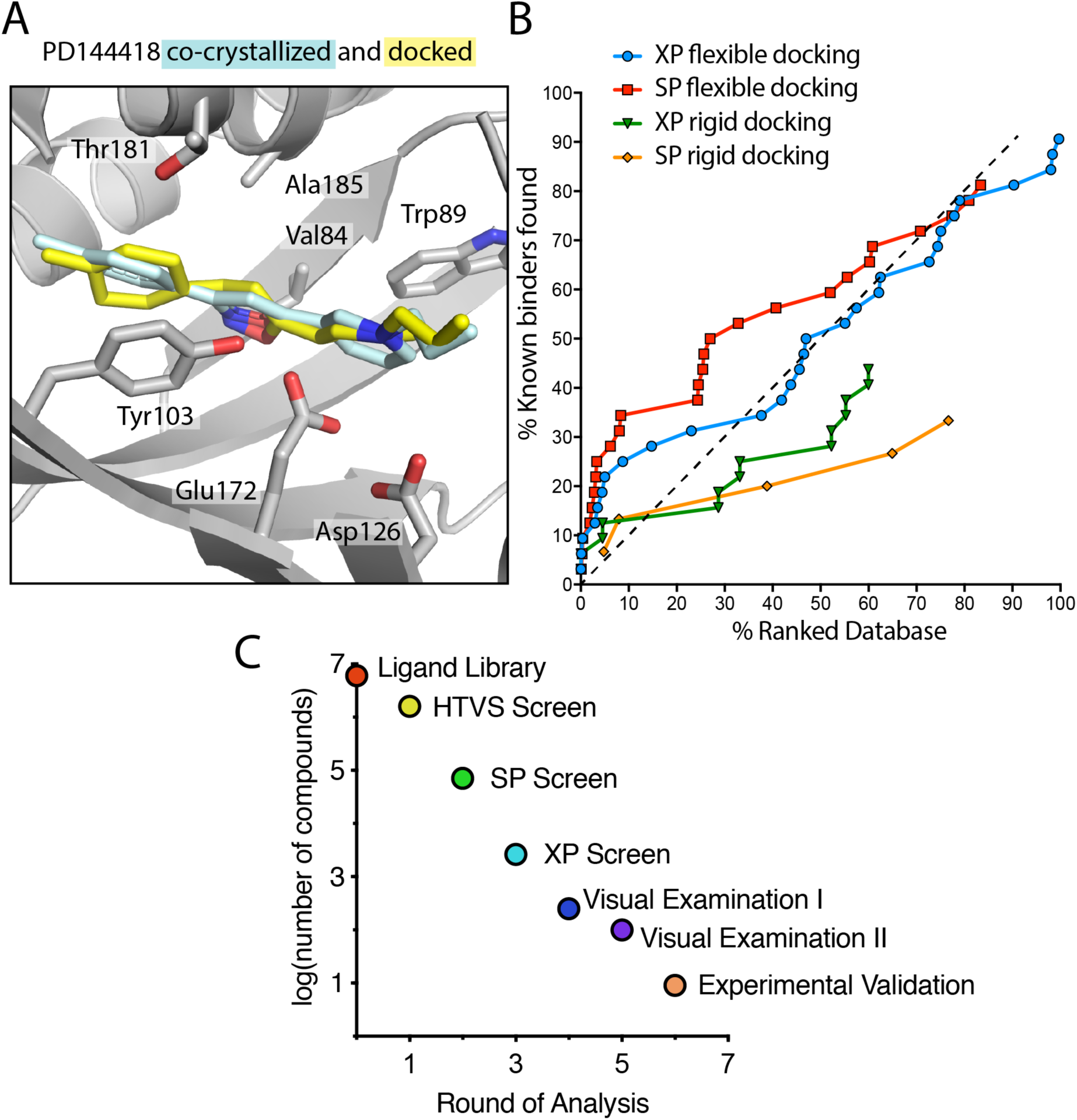
Ligands can be accurately docked into the σ_1_ receptor structure. (A) Superposition of the structures of PD144418 from the σ1 receptor crystal structure (cyan, PDB: 5HK1) and the highest-ranked docked pose (yellow). (B) Enrichment Plot showing the number of ligands recovered by the docking algorithm in the ranked database as a function of number of ligands sampled with the grid centered between Glu172 and Asp126. Docking types shown are XP flexible docking (blue circles), SP flexible docking (red squares), XP rigid docking (green triangles), and SP rigid docking (orange diamonds). The black line of slope 1 indicates performance expected via random selection of compounds by the docking algorithm. (C) Log-scale plot of the number of compounds remaining in each round of the docking pipeline.

Superimposition of the enrichment plots generated at the varying precision levels demonstrated consistently higher recovery rates of known binders in flexible docking compared to rigid docking. Extra precision flexible docking yielded the highest recovery rate (number of known binders scored/total number of binders in subset tested) of known binders at 90.63%, followed by standard precision flexible docking at just under 81.25%, extra precision standard docking at 43.75%, and standard precision rigid docking at just below 33.33%. However, flexible SP docking appeared better at recovering known binders early in the ranked database. It is interesting that SP recovered more known binders quickly, as XP is considered to be the most rigorous scoring paradigm, thus it would have been reasonable to see XP recover more known binders overall and recover more in the top 10% of the tested ligands. The relationship between SP and XP flexible and rigid docking is shown in Figure 2B.

In the docking experiment, compounds were selected from a starting library of over 6 million compounds through four sequential iterations of the docking protocol with increasing precision levels. Approximately1.6 million compounds passed through high-throughput virtual screening (HTVS) and went into SP, 71189 compounds from SP to XP, and 2625 compounds from XP to Flexible XP docking (see Figure 2C). From HTVS to SP all ligands that scored better than the re-docked co-crystallized ligand were selected. For the transition from SP to XP, the mean of all output scores in SP was found and all scores beyond 2 standard deviations from that mean were selected, and for all other precision levels chose ligands further than 1 standard deviation from the mean.

K-means clustering was applied to ligands based on volume occupied in binding site in an attempt to select a unique set of ligands to experimentally validate. After collapsing many sparse clusters, we found 3 main clusters with a total of 250 chemically and structurally diverse compounds. After visual inspection of the docked structures, the top 100 compounds were ranked based on glide score and the chemical plausibility of the docked pose. Of these, 17 representative compounds that were chemically divergent from one another and significantly different from existing σ1 receptor ligands were chosen for experimental assessment.

Among compounds purchased, 9 passed LC/MS inspection for purity and expected molecular weight and were characterized further (Figures S1 and S2). Figure S3 depicts the results from our initial screen of the compounds. Of the 9 compounds experimentally tested, 7 bound the σ1 receptor with <1 μM affinity (Table 1 and Figure S3B). Compound **2** bound with 10 nM affinity (Table 1). Compounds **3, 4, 5** and **9** also bound with < 100 nM affinity, while compounds **6** and **8** bound with 132 and 550 nM affinity respectively.

**Table 1.**
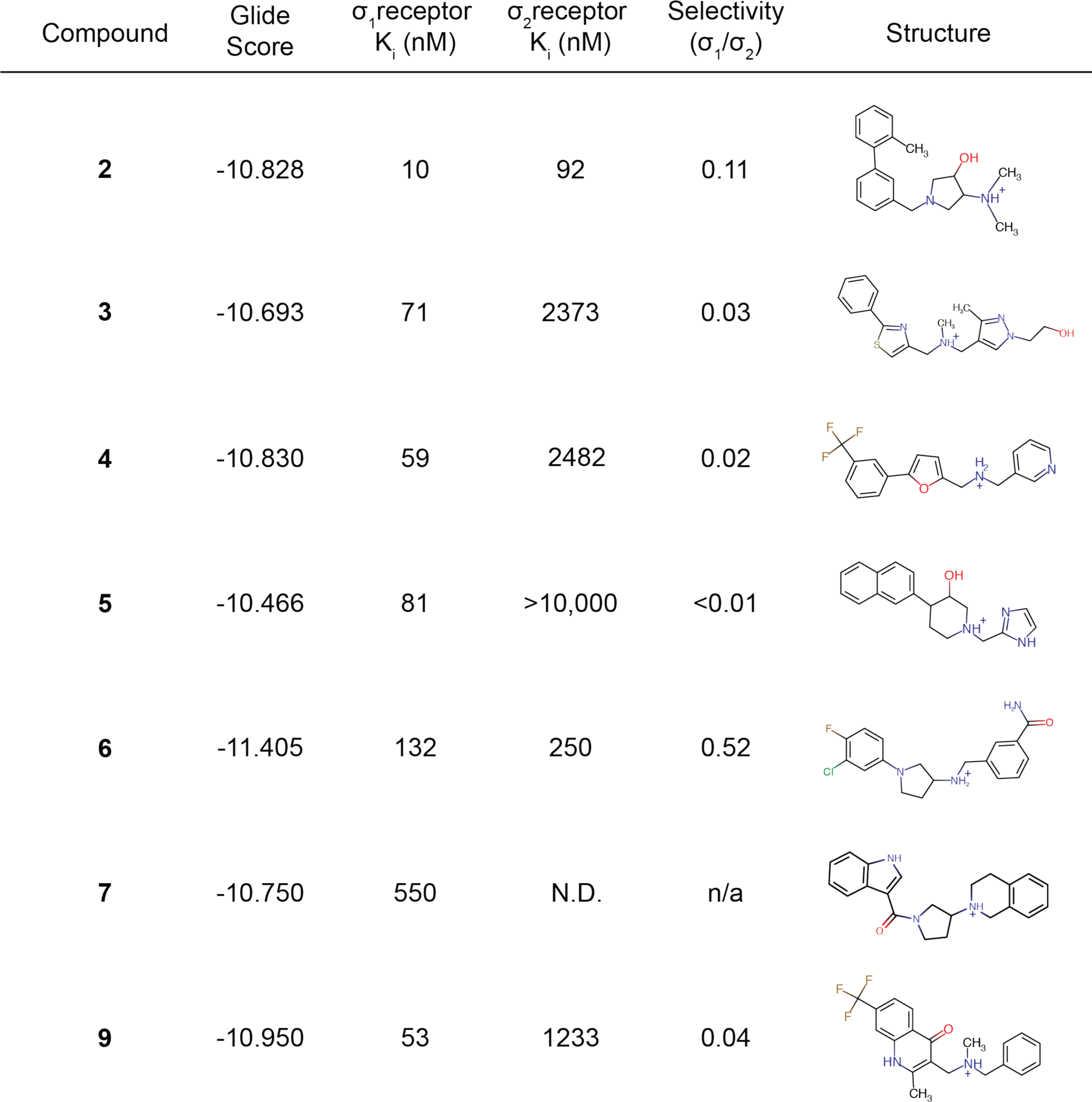
Summary of experimental characterization of docked compounds. All affinity values are representative results from at least two independent experiments performed in triplicate. “N.D.” indicates that the affinity was not determined.

| Compound | Glide Score | $\sigma_1$receptor<br>$K_i$ (nM) | $\sigma_2$receptor<br>$K_i$ (nM) | Selectivity<br>($\sigma_1/\sigma_2$) | Structure |
| --- | --- | --- | --- | --- | --- |
| 2 | -10.828 | 10 | 92 | 0.11 |  |
| 3 | -10.693 | 71 | 2373 | 0.03 |  |
| 4 | -10.830 | 59 | 2482 | 0.02 |  |
| 5 | -10.466 | 81 | >10,000 | <0.01 |  |
| 6 | -11.405 | 132 | 250 | 0.52 |  |
| 7 | -10.750 | 550 | N.D. | n/a |  |
| 9 | -10.950 | 53 | 1233 | 0.04 |  |

Though the σ_1_ and σ_2_ receptors are genetically unrelated, they have very similar pharmacological profiles and development of selective compounds can be challenging^22^. We reasoned that our structure-guided approach and the chemical diversity of our ligands might allow the discovery of subtype-selective probes. The 9 compounds tested for σ_1_ receptor binding were also tested for cross-reactivity with the σ_2_ receptor (Figures S3C and S3D, Table 1). Of the compounds that bound well to σ_1_ receptor, only compound **2** bound σ _2_ with <100 nM affinity (Table 1). The other compounds did not bind as well to σ_2_ in the initial screen as they did to σ_1_, with most exhibiting >1 μM affinity for the σ _2_ receptor.

To our knowledge, this work represents the first purely structure-based virtual screen on the σ_1_ receptor without the use of additional pharmacophore modeling. We found several ligands that bound with high affinity and had good selectivity for the σ_1_ receptor over the pharmacologically similar σ _2_ receptor. This work demonstrates that the σ_1_ receptor is highly amenable to *in silico* structure-based drug discovery, and that such approaches represent a viable way to identify novel σ_1_ receptor ligands at relatively low cost.

The σ_1_ receptor is an especially interesting target for virtual screening because of its potential therapeutic relevance and the lack of functional assays suited to experimental high-throughput screening. Previous work has identified the minimal chemical features of σ_1_ ligands to be a basic amine flanked by two hydrophobic regions, one that is 6-10 Å and another that is 2.5-3.9 Å^23^. Later work has refined this model, but the basic elements have remained unchanged^24-24^.

There have been previous attempts to design and discover σ_1_ receptor ligands through computational pharmacophore modeling^24,25^. These studies found high-affinity binders but were limited to the discovery of compounds with very similar chemical features to reference ligands. In another effort, a computationally predicted structural model of the σ_1_ receptor was developed and used for ligand screening^28^. However, this model showed little similarity to the experimentally determined σ_1_ receptor structure, and it only led to the identification of compounds with poorer affinity than that of a reference ligand, indicating that it offered little if any value in guiding ligand discovery.

More recent efforts have combined structural information from the σ_1_ receptor crystal structure with pharmacophore modelling or quantitative structure-affinity relationship (QSAR) approaches^26,27^. One recent report used the crystal structure of the σ_1_ receptor bound to PD144418 (PDB ID: 5HK1) to align 180 σ_1_ receptor antagonists and develop a 3D-QSAR model for σ_1_ antagonists^26^. This model was able to accurately predict ligands that would bind the σ_1_ receptor with high affinity from the Drugbank Database of FDA approved drugs, with the top hits binding the receptor with 58 nM and 160 nM affinity^26^. Similarly, another recent study used the same crystal structure to develop a predictive pharmacophore model, which was able to accurately predict binders from a set of 3,707,672 conformers of 25,676 compounds previously screened for σ_1_ receptor affinity^27^. Interestingly, this study found that the pharmacophore model derived from the σ_1_ crystal structure was more accurate for this set of compounds than docking into the structure directly using LibDock^27,29,30^. Neither of these models were used to identify uncharacterized compounds, though some of the FDA approved drugs were not known to bind σ_1_ receptor previously^26,27^.

Here, we show that Schrödinger Glide docking allows for the identification of novel σ_1_ receptor ligands. Starting with a computational library of over 6 million compounds, we were able to select only 9 for experimental characterization and still found 7 binders with sub-micromolar affinity, corresponding to a success rate of approximately 77%. This compares favorably to other docking studies. Of 54 recent docking studies, only 8 had higher hit rates and only 3 found compounds with better affinity than the best hit of 10 nM reported here, despite the use of significantly larger screening libraries in some cases^31^. This may be because the σ_1_ receptor is particularly well suited to computational docking. Most high-affinity ligands found to date bind with only a single electrostatic interaction between a primary amine and Glu172, and affinity is likely driven largely by hydrophobic interactions and steric constraints. This relatively simple binding mode may be easier to accurately predict by docking than more complex ones. Indeed, constraining ligands to be docked near Glu172 and Asp126 was sufficient to successfully find binders, without requiring a more complex pharmacophore model.

In addition to having high affinity for the σ_1_ receptor, most compounds tested here exhibited high selectivity for the σ_1_ receptor over the pharmacologically similar σ_2_ receptor. Few compounds bound to σ_2_ with sub-micromolar affinity, and almost all tested compounds had higher affinity for the σ_1_ receptor than the σ_2_ receptor. For example, Compound **5** exhibited 81 nM affinity for the σ_1_ receptor yet bound with >10 μM affinity to the σ_2_ receptor. In contrast, compounds 2 and 6 showed high affinity for both σ_1_ and the σ_2_ receptor, indicating that non-selective ligands can also be identified by virtual screening even against just one receptor type. While we focused here on the σ_1_ receptor, relatively few highly σ_2_ selective compounds are currently available. The σ _2_ structure may allow identification of highly σ_2_-selective ligands using similar approaches to those described here, further expanding the suite of molecular tools available for σ receptor research.

## Materials and Methods

### Computational Methods

Prior to docking, the σ_1_ receptor structure (PDB ID: 5HK1) was put through the Protein Preparation Protocol^32^. This added hydrogens, removed co-crystallized water molecules, enumerated bond orders, optimized the receptor’s hydrogen bond network, and ran a restrained minimization to alleviate backbone clashes within the receptor backbone. Additionally, Asp126 was protonated in our docking screens on the basis of its apparent hydrogen bond with Glu172 in the crystal structure of the receptor^11^.

The eMolecules ligand library was prepared using the Ligand Preparation Protocol^32^ to generate three-dimensional geometries, assign proper bond orders, and generate accessible tautomer and ionization states for the ligands prior to virtual screening^33^.

The grid generation protocol in Glide Docking defines the target area (binding site) for each ligand to be tested within during the docking program. This was centered between the carboxylates of Glu172 and Asp126 and was kept constant as the docking precision was increased. Docking began with high throughput virtual screening (HTVS), in which all compounds in the eMolecules library were tested. All ligands with Glide scores more favorable than the score of the re-docked co-crystallized ligand PD144418 were put through standard precision screening (SP). All compounds scoring better than 1 standard deviation from the mean were tested in extra precision (XP), and the top 10% of XP ligands were put through flexible XP docking, where the receptor backbone was given freedom to shift in order to fit ligands to the orthosteric site.

To preform enrichment analysis similar to previous approaches^34^, a set of 15 known σ_1_ receptor binders were collected from the PDSP Ki Database^35^ and put through the ligand preparation protocol. These ligands were then tested in varying precision levels after being merged with a set of 100,000 eMolecules ligands of similar weight to the known binders. The average molecular weight of the known binders taken from the PDSP Ki Database is 351.07 g/mol, and the sample of ligands that were selected for enrichment with known binders had molecular weights between 300 and 400 g/mol in an attempt to force Glide to focus on differences in chemical composition rather than physicochemical descriptors.

Schrodinger Drug Discovery Platform installation was supported by SBGrid Consortium and all data was generated and processed on SBGrid servers.^20^

### Recombinant receptor expression and preparation of *Sf*9 cell membranes

Membranes were prepared from *Sf*9 insect cells infected with baculovirus encoding either recombinant σ_1_ or σ_2_ receptor, using a protocol adapted from that of Vilner et al., as previously described^10,11,22^. Briefly, either the σ_1_ receptor or σ_2_ receptor, which was recently identified as TMEM97^10^, were expressed in Sf9 cells using the pFastBac baculovirus expression system. Cells were lysed by osmotic shock and membrane fractions were separated by repeated centrifugation and washing steps, followed by dounce homogenization. Total protein content was quantified using the BioRad DC protein assay, and membranes were stored in 100 μL aliquots at a concentration of 3-10 mg/mL at −80 °C until use.

### Screening docking hits for σ_1_ and σ_2_ receptor binding

Membrane radioligand binding assays were performed as described with slight modifications^10,11,36^. All compounds were diluted to 10 μM in 50 mM Tris pH 8.0. Membrane samples were also diluted and homogenized in 50 mM Tris pH 8.0 using a needle and syringe. A total of 2.5 μg of protein was added to each well of a 96-well block in a final volume of 100 μL, which also contained 1 μM of the compound and 10 nM of the appropriate radioligand (^3^H (+)-pentazocine for the σ_1_ receptor, and ^3^H DTG for the σ_2_ receptor). All reactions were performed in triplicate. Haloperidol was used as a positive control, and buffer alone served as a negative control. Samples were incubated at room temperature with shaking for 1.5 h. The reaction was then terminated and samples were then applied to glass fiber filters (Merck Millipore) using a Brandel cell harvester. Filters were soaked with 0.3% (w/v) polyethylimine prior to use. After harvesting, filters were soaked overnight in scintillation vials containing 5 mL of Cytoscint scintillation fluid and measured on a Beckman Coulter LS 6500 scintillation counter.

### Saturation binding in *Sf*9 cell membranes

^3^H (+)-pentazocine and ^3^H DTG saturation binding to *Sf*9 membranes respectively expressing σ_1_ or σ_2_ receptor were determined using a saturation binding assay similar to that described by Chu and Ruoho^36^,which we have described previously^10,11^. The membranes (2.5 μg and 0.5 μg total protein per reaction for σ_1_ and σ_2_ expressing membranes, respectively) were incubated in a 100 μL reaction buffered with 50 mM Tris pH 8.0, containing 0-300 nM ^3^H (+)-pentazocine for σ_1_ binding, or 0-300 nM ^3^H DTG for σ_2_ receptor binding. In the σ_2_ receptor binding assay, concentrations of 100 and 300 nM DTG were achieved by isotopic dilution to minimize ^3^H DTG expenditure. For both receptors, nonspecific binding was assayed by parallel reactions containing 2 μM haloperidol. Reactions were incubated at 37 °C for 1.5 h and then terminated, filtered, and measured as for the single-point binding described previously. All reactions were performed in triplicate in a 96-well block. *K*D values were calculated using non-linear regression tools in Graphpad Prism.

### Competition binding assays in *Sf*9 membranes

^3^H (+)-pentazocine or ^3^H DTG competition curves testing the binding the docked compounds to the σ_1_ and σ_2_ receptors respectively, were performed similarly to the protocol described by Chu and Ruoho^36^,with slight modifications we have used previously^10,11^. Briefly, *Sf9* insect membranes overexpressing either σ_1_ receptor (2.5 μg of total protein per reaction) or σ_2_ receptor (0.5 --2.5 μg total protein per reaction) were incubated in a 100 μL reaction buffered with 50 mM Tris pH 8.0, with the appropriate radioligand at a 10 nM concentration and eight competing cold compound concentrations ranging from 10 pM – 100 μM. Reactions were incubated at 37 °C for 1.5 h to reach equilibrium, and then terminated by filtration as described previously. *K*_*i*_ values were computed by directly fitting the data and using the experimentally determined probe K_D_ to calculate a K_i_ value, using the Graphpad Prism software. This process implicitly uses a Cheng-Prusoff correction, so no secondary correction was applied.

### LC/MS analysis of compound purity

Purchased compounds were analyzed on an Agilent QTOF 6530. Samples were separated by reverse-phase HPLC (Agilent Extend C18 column 1.8 μM, 50 × 2.1 mm) at a flow rate of 0.3 mL min^-1^ in H_2_O with 0.1% formic acid for 15 min followed by a 5 min elution in 10-100% acetonitrile. Data were analyzed using Agilent MassHunter Quantitative Analysis Software.

HRMS (ESI+) *m/z* calcd for C_20_H_31_N_3_O [M + H] 342.2540, found 342.2545 (error 1.5 ppm);

HRMS (ESI+) *m/z* calcd for C_20_H_26_N_2_O [M + H] 311.2118, found 311.2117 (error 0.3 ppm);

HRMS (ESI+) *m/z* calcd for C_18_H_22_N_4_OS [M + H] 343.1587, found 343.1587 (error <0.3 ppm);

HRMS (ESI+) *m/z* calcd for C_18_H_15_N_2_OF_3_ [M + H] 333.1209, found 333.1231 (error 6.6 ppm);

HRMS (ESI+) *m/z* calcd for C_19_H_21_N_3_O [M + H] 308.1757, found 308.1764 (error 2.3 ppm);

HRMS (ESI+) *m/z* calcd for C_18_H_19_N_3_OClF [M + H] 348.1273, found 348.1279 (error 1.7 ppm);

HRMS (ESI+) *m/z* calcd for C_22_H_23_N_3_O [M + H] 346.1914, found 346.1912 (error 0.6 ppm);

HRMS (ESI+) *m/z* calcd for C_19_H_19_N_2_OF_3_ [M + H] 349.1522, found 349.1538 (error 4.6 ppm);

HRMS (ESI+) *m/z* calcd for C_20_H_19_N_2_OF_3_ [M + H] 361.1522, found 361.1545 (error 6.4 ppm).

## Author Information

Supporting Information: Binding data, Compounds tested in binding assays, eMolecules ligand library information, Compound similarity

## Author Contributions

D.A.G. designed and carried out computational docking and analysis of results, and together with H.R.S. performed radioligand binding assays and associated data analysis. H.R.S. conducted LC-MS for assessment of compound purity. P.S. Co-designed the computational pipeline and co-supervised the docking experiments. A.C.K. supervised the work and together with D.A.G. and H.R.S. wrote the manuscript.

## Acknowledgements

We thank Dr. Matthew Henke and the Harvard Medical School Analytical Chemistry core, and Dr. Meredith Skiba for assistance with LC/MS. This work was supported by NIH grant R01GM119185 (A.C.K.), and grant number FG-2017-9226 from the Alfred P. Sloan Foundation (A.C.K.). H.R.S is supported by an NSF Graduate Research Fellowship DGE1745303. D.A.G is supported by BCMP Scholars Internship Program and Molecular Biophysics Training Grant NIGMS T32 GM008313.

## Abbreviations Used

NMDA: N-methyl-D-aspartate
ECFP: Extended Connectivity Fingerprints
HTVS: High Throughput Virtual Screening
SP: Standard Precision Virtual Screening
XP: Extra Precision Virtual Screening
4-IBP: N-(benzylpiperidin-4yl)-4-iodobenzamide
PD144418: 1,2,3,6-tetrahydro-5-[3-(4-methylphenyl)-5-isoxazolyl]-1-propylpyridine
NE-100: *N, N*-dipropyl-2-[4-methoxy-3-(2-phenylethoxy)phenyl]-ehthylamine monohydrochloride
DTG: 1,3-Di-(2-tolyl)guanidine

