## Supplementary Information for "Virtual screening identifies novel high-affinity σ_1_ receptor ligands" (from another version of this preprint: version 1, 2019-07-11)

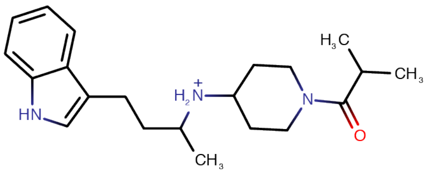

1

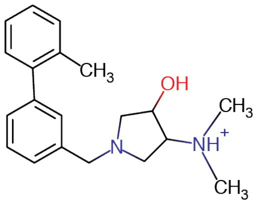

2

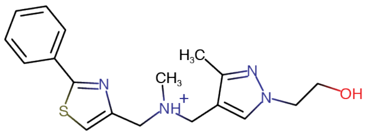

3

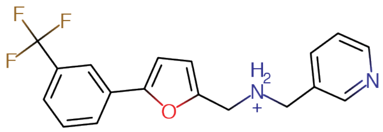

4

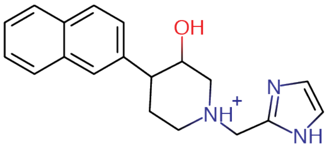

5

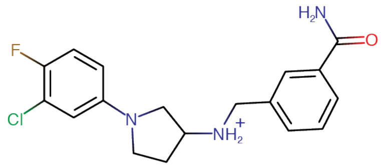

6

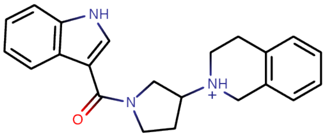

7

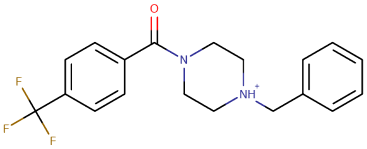

8

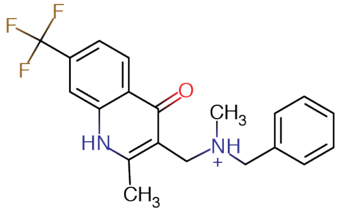

9

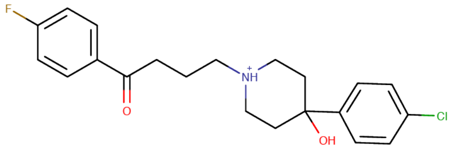

Haloperidol (control ligand)

**Figure S1.** Chemical structures of the ligands tested in $\sigma$ receptor binding assays.

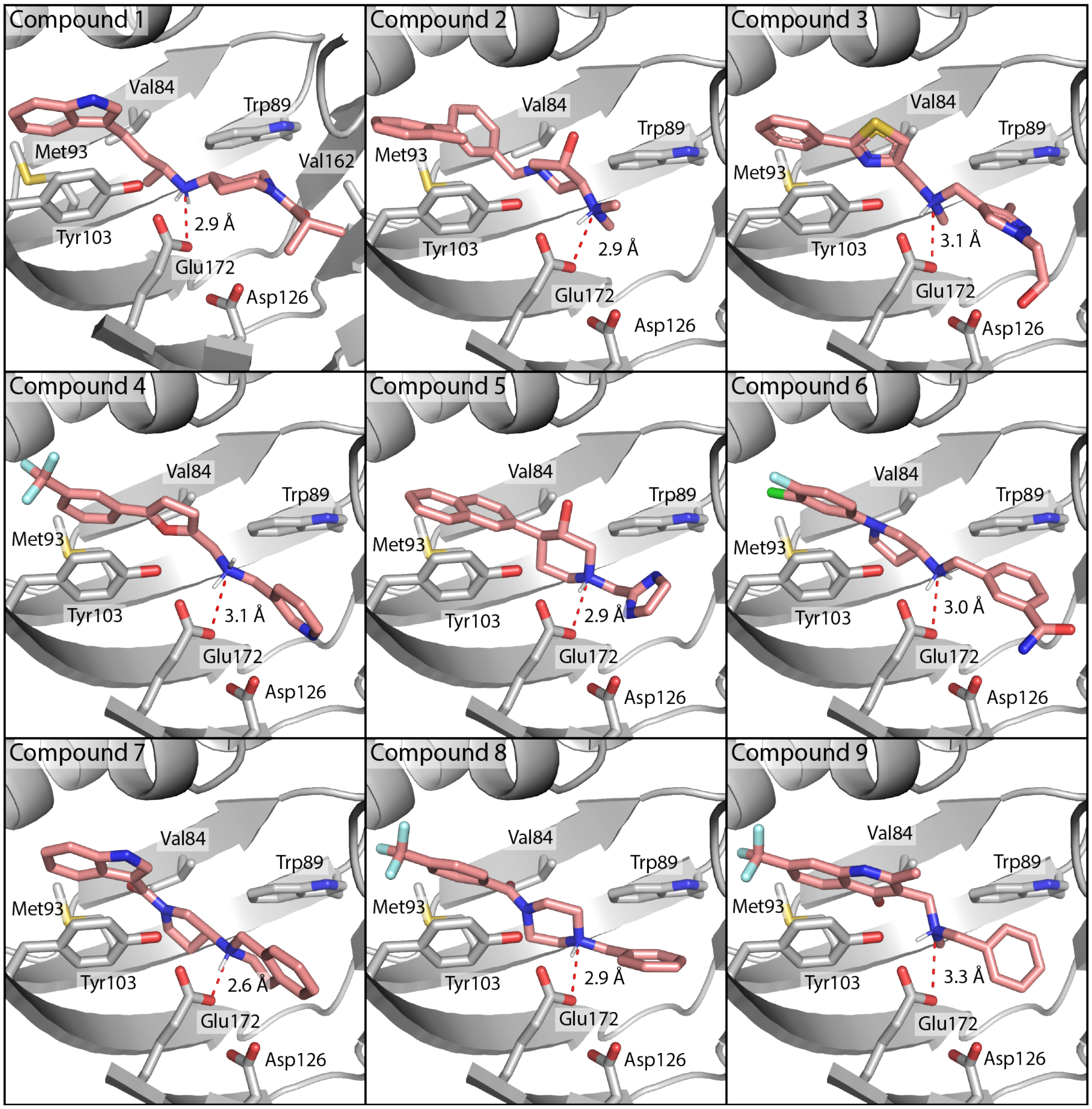


**Figure S2.** Docked poses of the nine compounds chosen for experimental characterization. The $\sigma_1$ receptor is shown in grey, and the ligands are shown in pink.

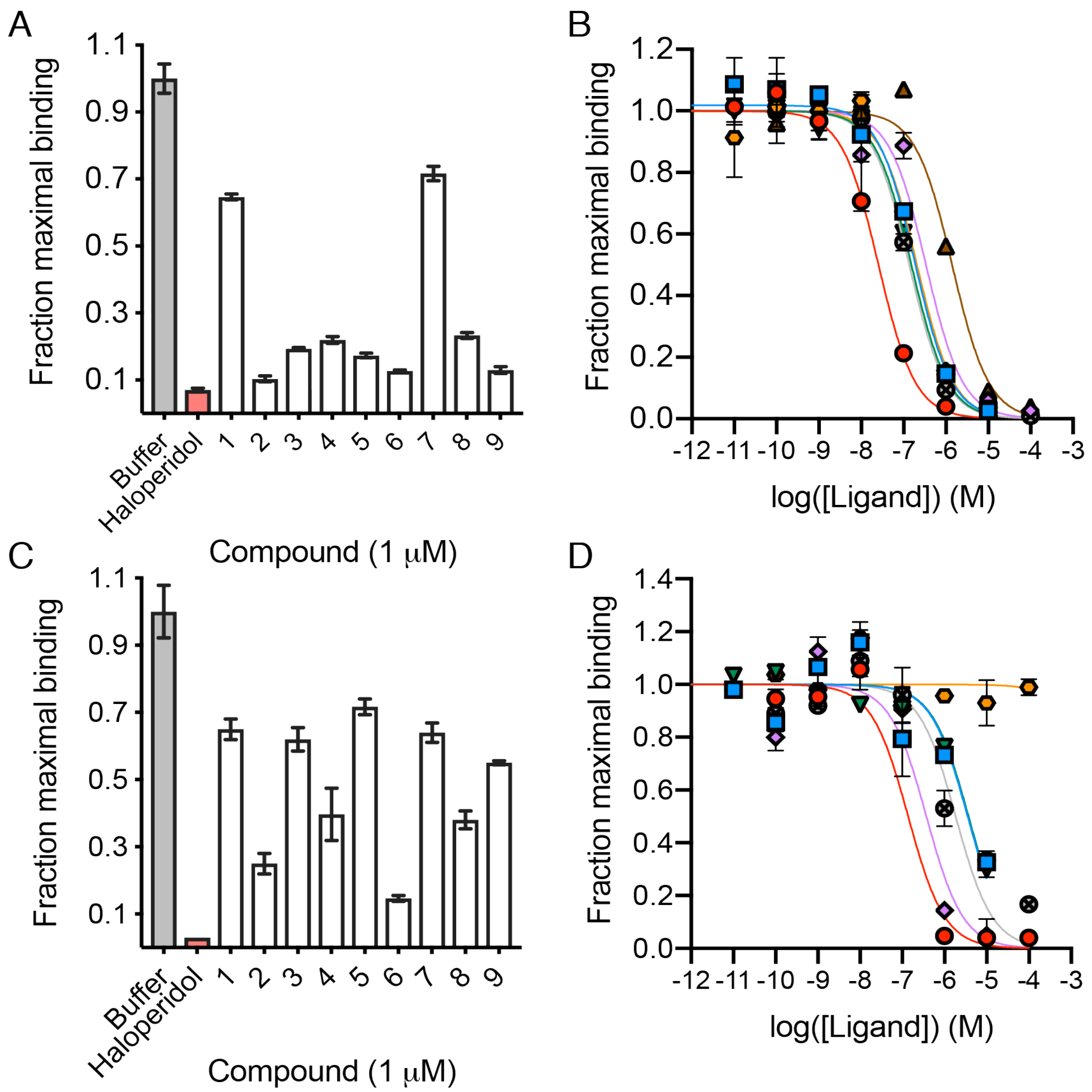


**Figure S3.** Experimental characterization of compounds **1**-**9**. All measurements were taken in triplicate. Error bars represent standard error of the mean. (A) Single-point competition binding assay of each compound at 1 μM against 10 nM [$^3$H](+)-PTZ. (B) Competition binding curves of compounds **2** (red circles), **3** (blue squares), **4** (green triangles pointing down), **5** (orange hexagons), **6** (lavender diamonds), **7** (brown triangles pointing up), and **9** (grey circles with a black "x") against 10 nM [$^3$H](+)-PTZ. Curves are representative of at least two independent experiments performed in triplicate. (C) Single-point competition binding assay of each compound at 1 μM against 10 nM [$^3$H]DTG. (D) Competition curves against 10 nM [$^3$H]DTG. Colors are the same as in (C). Curves are representative of at least two independent experiments performed in triplicate.
